# Old growth attributes by chain saw: how between-patch heterogeneity changes the metacommunities of beetles in temperate forests

**DOI:** 10.1101/2025.08.30.673209

**Authors:** Oliver Mitesser, Marc Cadotte, Akira S Mori, Fons van der Plas, Anne Chao, Julia Rothacher, Claus Bässler, Mirjana Bevanda, Peter H. W. Biedermann, Pia Bradler, Antonio Castaneda-Gomez, Orsi Decker, Benjamin M. Delory, Sebastian Dittrich, Heike Feldhaar, Andreas Fichtner, Alexander Kreis, Lisa Köstler-Albert, Ludwig Lettenmaier, Goddert von Oheimb, Luise Pflumm, Kerstin Pierick, Jakob Schwalb-Willmann, Simon Thorn, Leah Vogelfänger, Wolfgang Weisser, Martin Wegmann, Clara Wild, Jörg Müller

## Abstract

Metacommunity theory has expanded our understanding of how spatial dynamics and local interactions influence species communities. Different assembly archetypes, reflecting different roles of species differences, habitat differences, and dispersal have been described, but we lack empirical studies specifically in terrestrial habitats testing which archetype is most important. In a replicated design we experimentally enhanced structural between-patch heterogeneity in homogeneous production forests and developed a statistical framework controlling for sample incompleteness to detect different metacommunity processes. Meta-analyses on >100K individuals of >1.3K beetle species showed an increase of ∼60 species in heterogenized forests at γ-level promoted by increasing α-diversity consistent with the *mass-effect* and an increase of β-diversity by ∼10% supporting *species-sorting*. Additionally, we tested β-deviations from random assembly as a proxy of *neutral processes*. Findings indicate that enhancing structural heterogeneity can shift forests from *patch-dynamics* dominance towards *mass-effect* and *species-sorting*, offering a promising pathway to restore biodiversity in managed landscapes.

## Introduction

During the last two decades, metacommunity theory has expanded our understanding of how spatial dynamics and local interactions influence the structure and diversity of species communities (Leibold *et al*. 2004; Cottenie 2005; Gonzalez 2009; Record *et al*. 2021). Within this framework, four archetypes have been developed to characterize metacommunities (Logue *et al*. 2011; Leibold & Chase 2018). Each of them predicts different assembly mechanisms based on dispersal and differences in habitats and species: The *patch-dynamics* archetype assumes homogeneous patches which are occupied by species with a trade-off between dispersal and local dominance. Superior competitors can displace others in occupied patches, whereas weaker competitors may persist at the regional level by rapidly colonizing newly available patches. Variation in species composition is due to processes of colonization and extinction. The *species-sorting* archetype assumes that habitat patches are heterogeneous, and dispersal capacity is just sufficiently high to reach the different patches. Local community composition is primarily shaped by a match between environmental conditions and the ecological niches of the species. A high level of environmental variation across the landscape thus leads to high regional species richness through different species assemblages in different patches. The *mass-effect* archetype assumes that heterogeneous habitat patches are interconnected, and species’ dispersal abilities are high. This allows species to reproduce in source patches and frequent dispersal allows them to also exist in sink patches where local conditions prevent long-term survival or reproduction. Massive dispersal over long distances might override signals of species-niche matching in the overall biodiversity response. Last, the *neutral* archetype relies on the assumption that individuals of different species are equivalent in survival, reproduction, and dispersal abilities. Assembly processes entirely depend on stochastic events including immigration and extinction (Hubbell 2001). Ecological drift and rare speciation shape regional diversity patterns over large time scales and large metacommunities serve as persistent sources of species for local communities.

Theoretical studies have shown that the different metacommunity archetypes make different predictions on how biodiversity is distributed across space and which predictors explain the patterns (Thompson *et al*. 2020). While processes of all archetypes have been demonstrated to matter, these relationships can be utilized to identify the most influential mechanisms in real-world landscapes (see hypotheses below). Despite the theoretical progress in metacommunity theory, it is still insufficiently linked to empirical data, particularly in terrestrial habitats (Logue *et al*. 2011), where soft boundaries between patches with gradual transitions of resources, community composition, and processes are the rule rather than the exception. To disentangle the importance of different metacommunity archetypes, Logue *et al*. (2011) proposed developing an integrative framework that combines new experimental approaches and novel statistical methods and is based on three key axes: species differences (functional equivalence vs. differentiation), habitat differences (homogeneity vs. heterogeneity), and dispersal dynamics (Fig. 1a).

**Figure 1:**
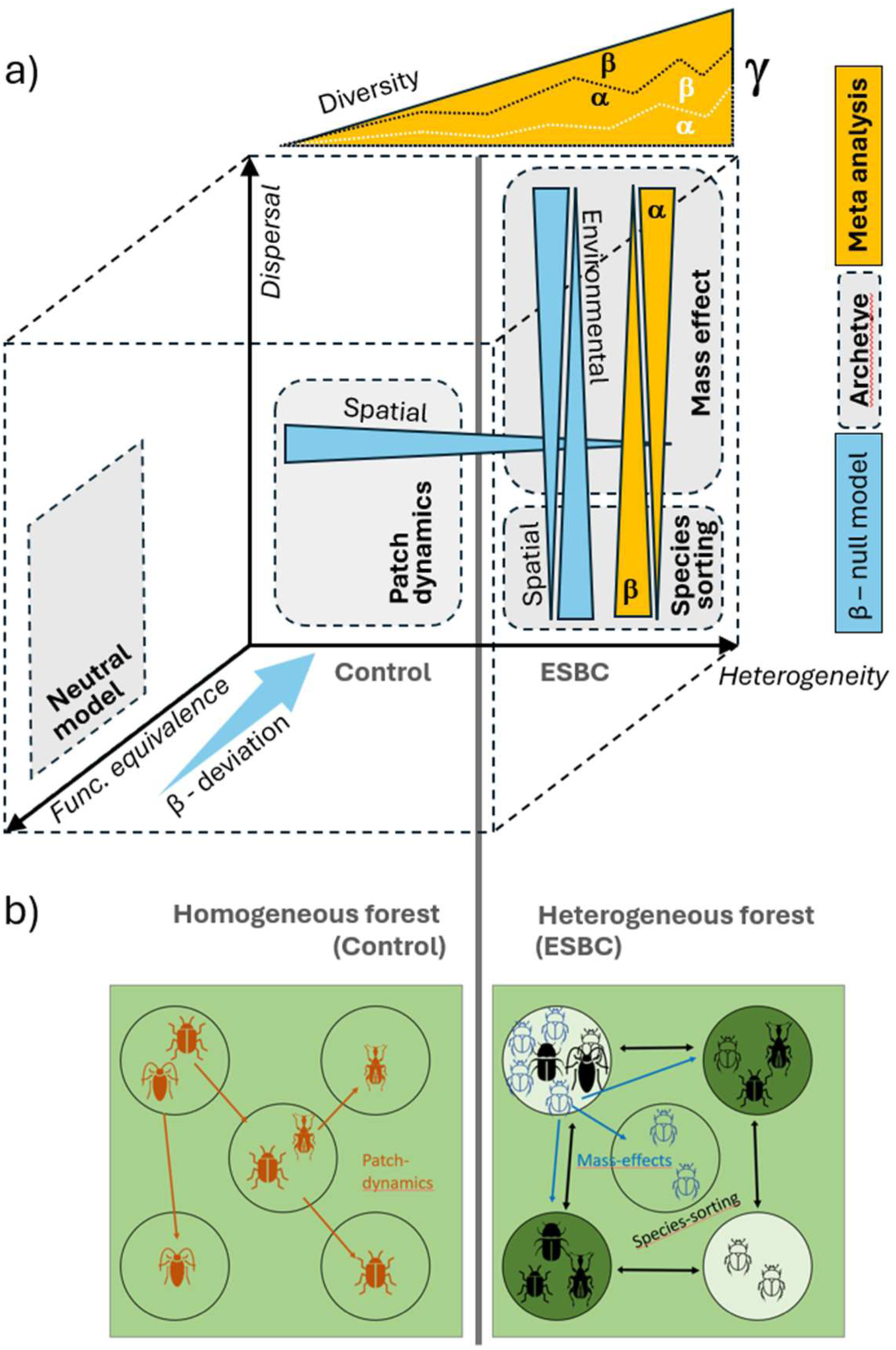
a) Extension of Logue et al. (2011)’s scheme of metacommunity mechanisms based on the dimensions of heterogeneity, functional equivalence of individuals, and dispersal. Metacommunity archetypes of *species-sorting*, *mass-effect*, *patch-dynamics*, and *neutral processes* occupy typical areas in the diagram. Both meta-analyses (yellow) and β-deviation analysis (blue) allow discrimination between the processes. Dotted black and white lines within the hypothetical increase of γ-diversity (yellow triangle at the top) indicate different hypothetical allocation of the effect to α- and β-diversity. b) Conceptual figure on archetypes from metacommunity theory explaining diversity and assembly mechanisms in local beetle communities in homogeneous and heterogenous forests. White and green colours indicate habitat differences. In ESBC patches light and dark beetle symbols indicate matching of beetles’ attributes to light and dark green habitats, respectively.

We selected homogeneous production forests as a model system to implement a novel large-scale experimental approach (Müller *et al*. 2023) aimed at identifying the dominant processes driving metacommunity assembly and assessing how their relative importance varies with landscape heterogeneity. At the beginning of the experiment, we manipulated the between-forest-patch heterogeneity by creating gaps and deadwood towards old-growth features by chainsaw and selected homogeneous control forests. In parallel, we developed a new meta-analysis approach to disentangle the effects of within-patch α- and between-patch β-diversity on γ-diversity at landscape scale while accounting for sample incompleteness in empirical data for taxonomic (TD), functional (FD), and phylogenetic diversity (PD) along Hill numbers to capture differences in species abundances and ecological attributes (Chao *et al*. 2023). Considering these multiple dimensions of diversity (facet: TD, FD, PD; scale: α, β, γ; Hill number: q=0, 1, 2) enables the identification of underlying processes structuring species assemblages, depending on whether diversity is shaped by distinct functional traits, local versus regional patterns, or rare versus dominant species, respectively (Jeliazkov & Chase 2024).

The design allowed us to use the generalized concept of diversity decomposition efficiently in a replicated experiment at the landscape level. In the same framework, we were able to estimate pairwise β-deviations (from randomized null models) between-patches of a forest to create null-model communities (*neutral archetype;* Logue *et al*. 2011) avoiding well documented artefacts in β-diversity by under-sampling (Tuomisto 2010; Tucker *et al*. 2016). Subtraction of null-model β-diversity in general controls for effects by demographic and dispersal stochasticity on the β-diversity pattern expected from the *neutral* archetype (Tucker *et al*. 2016). β-deviation and spatial and environmental distances offered the option of deciphering the assembly mechanisms of metacommunities.

We used beetles as one of the taxonomically, phylogenetically, and functionally most diverse insect orders (Farrell 1998) which are particularly vulnerable under climate change (Thom & Seidl 2016). We expected a shift from metacommunities dominated by *patch-dynamics* in homogeneous control forests to metacommunities structured by *species-sorting* and *mass-effects* stimulating our hypotheses about diversities (α, β, and γ) and β-deviation. We first test the hypothesis (H1) that enhancement of between-patch heterogeneity increases γ-diversity of a forest. Along with this hypothesis, we predict an increase of γ-diversity by an increase of β-diversity according to *species-sorting* or by an increase of α-diversity by the *mass-effect* promoting source patches by resource enrichment, but at the same time eventually reducing β-diversity (Mouquet & Loreau 2003) (Fig. 1). Secondly, we hypothesise (H2) that β-deviation is better explained by spatial distance in homogeneous forests because other environmental attributes as potential alternative predictors have only low variation. In heterogeneous forests where *species-sorting* can increase the correlation between communities and environment we hypothesise (H3) environmental variation to be the better predictor of β-deviation. As species differences in habitat requirements and in mobility affect *species-sorting* and dispersal we expect effects of habitat difference in heterogeneous forests specifically on functional/phylogenetic β-deviation.

Our results demonstrate an increase in α-, β-, and γ-diversity in response to ESBC treatment and suggest that metacommunities in homogeneous forests are dominated by *patch-dynamics*, while those in heterogeneous forests by *mass-effect* and *species-sorting*.

## Materials and methods

### Study area and experimental design

We established our pairwise design of a district experimentally enhanced in structural beta complexity (ESBC) and a homogeneous control district at 11 sites throughout Germany covering a wide range of climate (c. 50 -1100 m a.s.l.) and soil conditions (acidic to calcareous) in European beech (*Fagus sylvatica*) mixed forests (Müller *et al*. 2023). In total, the study included 22 districts, and within each district a set of 9 or 15 patches was established, with a spatial extent of 50 m × 50 m in a randomized block design. The interventions aimed to enhance the variation in canopy density and deadwood structures. Manipulations were carried out in the winter of 2015/16 (Bavarian Forest, Passau), 2016/17 (Saarland, Lübeck, Hunsrück) and 2018/19 (University Forest, i.e. the forest of the University of Würzburg) to allow sufficient time for the colonization by forest fauna. LiDAR measurements validate that the ESBC districts remained structurally more heterogeneous than the control districts during the study period (Pierick *et al*. 2025, *preprint*).

**Figure 2:**
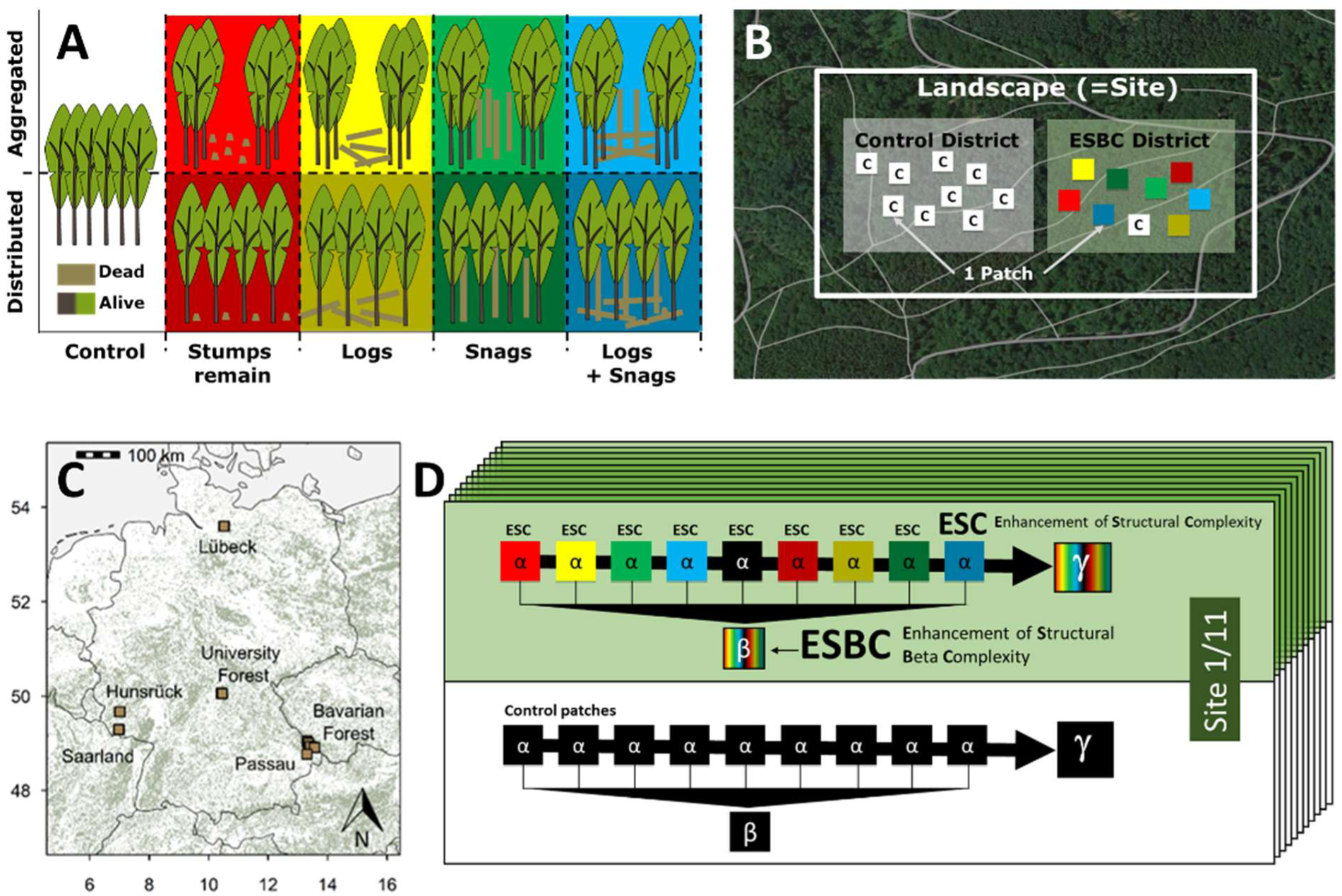
Design of replicated manipulation of habitat heterogeneity in 11 homogeneous production forests and their 11 control forests. A) Interventions were either aggregated in the centre of the patch (upper row) or evenly distributed over the patch area (lower row). B) Site with treatment (ESBC: Enhancement of Structural Beta Complexity) district and control district. C) Geographical distribution of the 11 sites. D) Schematic representation of the ecosystem elements site, district, and patch in relation to derived diversity levels for an exemplary site.

### Beetle data

In each patch we set up two flight interception traps and two pitfall traps to reach sufficient sample size (number of individuals) to represent the local environmental conditions (see (Müller & Brandl 2009). Sampling was conducted from May to July 2022 in the University Forest (3 sites), the Bavarian Forest (4 sites), and the Passau region (1 site), while the remaining sites were sampled in 2023. These months represent >90% of individuals in beetle sampling during the vegetation period. Traps were emptied monthly and sorted on order level. Species identification was conducted by professional taxonomists. As traits we selected 1) body size as a universal trait for aspects of reproduction, dispersal ability, and general population size (Chown & Gaston 2010), 2) brightness level as an important trait for species response to climate conditions (Zeuss *et al*. 2014), 3) the adoption of a life style in deadwood (saproxylic), and 4) the feeding strategy. All trait information was taken from literature (Freude *et al*. 1964-1983; Koch 1989-1992; Seibold *et al*. 2015; Hagge *et al*. 2021). As phylogenies of beetles are still incomplete, we used as a surrogate a phylogeny based on taxonomic classification (Freude *et al*. 1964-1983; Abrego *et al*. 2024).

### Environmental and spatial difference

To characterize the habitat dissimilarity of patches in addition to their spatial distance, we calculated the environmental distance between two patches of a district as their difference in quantitative attributes of forest structure. We used canopy densities measured by LiDAR via drone flights using the penetration ratio at 6 m above ground (Müller & Brandl 2009) and the volume of deadwood per patch to quantify the level of ESBC. Variation in both variables was predominantly driven by our experimental ESBC manipulation. However, as natural tree mortality is still increasing in Europe (Senf *et al*. 2018), both also capture add-on natural dieback processes. There might be additional more subtle variation or changes between forest patches in environmental conditions, e.g. soil moisture, not captured by LiDAR and deadwood volume. Therefore, we additionally calculated unweighted mean Ellenberg-indicator values based on plant relevés estimated according to a simplified percentage scale (Dittrich et al. 2013) and recorded in all patches (Ellenberg *et al*. 1991; Ewald 2003), which has been proven to be highly integrative in describing such subtle environmental conditions (Ewald 2003, Müller et al. 2009). We used the difference between patches of these values as a proxy for abiotic distance. To avoid overlap between information in structure measured by LiDAR and these indicator values we removed the light indicator value. Spatial distance between patches was calculated in meters based on projected coordinates in the ETRS1989 format.

## Statistical methods

### Meta-analyses

Statistical analyses were conducted in R (version 4.3.3) (R Core Team, 2023). To allow for direct comparison, taxonomic (TD), phylogenetic (PD), and functional diversity (FD) were assessed as the effective number of equally abundant species, divergent lineages, and distinct functional groups and for Hill numbers q = 0, 1, 2 focusing on rare, common, and dominant species. These facets of beetle diversity and corresponding confidence intervals were calculated for each of the 22 districts, utilizing the iNEXT.3D package (Chao *et al*. 2022) and the included bootstrapping approach with 200 replicates. γ-diversity (at the landscape level) was decomposed into α (patch level) and multiplicative β-diversity using iNEXT.beta3D (version 2.0.8) (Chao et al., 2023). We applied the 1-S transformation, a normalized dissimilarity measure equivalent to Jaccard-type turnover, mapping β-diversity to the range of 0 indicating identical species composition among patches to 1 representing completely disjunct communities. Standardization also controls for variation in patch numbers of sites (9 and 15) (Chao *et al*. 2019). For comparison between ESBC and control districts, we calculated differences in diversity metrics (TD, PD, and FD in combination with α-, β-, and γ-diversity, as well as with q = 0, 1, and 2) for each of the 11 district pairs with positive values indicating higher diversity in the ESBC district. Differences from the 11 forest pairs were weighted and combined by meta-analysis (R-package in preparation: https://github.com/AnneChao/iNEXT.meta) propagating district-wise bootstrapping confidence intervals to the aggregated metric and being presented as error bars in Fig. 3. Confidence intervals that exclude the value 0 indicate significant differences between ESBC and control districts. To control for variation in sampling effort eventually correlating with the treatment, we standardized diversity estimates to a common coverage level of 0.95 at the maximum of the observed distribution (Chao *et al*. 2020). District-specific results are shown in the Supplement, figures S1-S27.

**Figure 3:**
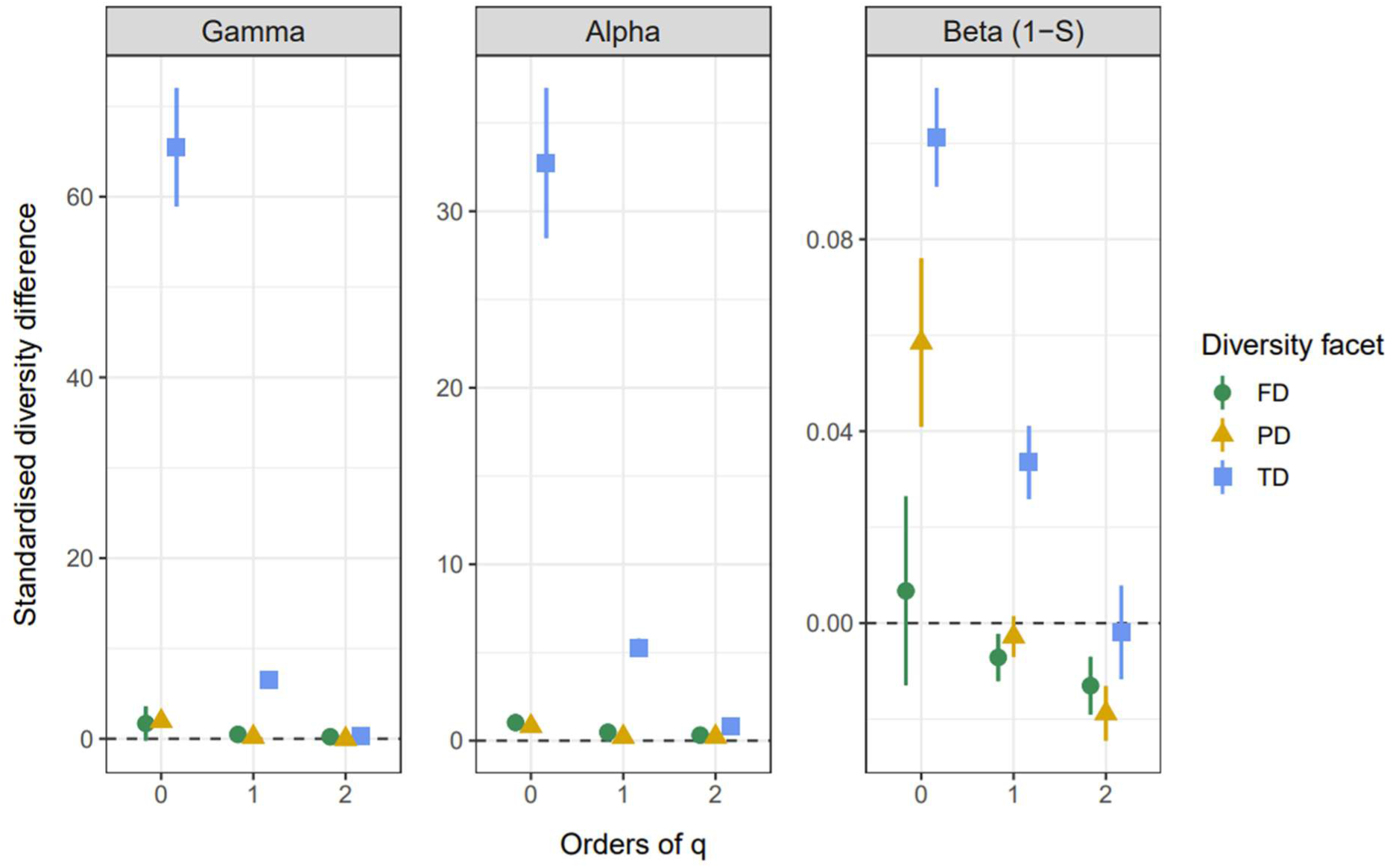
Results of the meta-analyses based on 11 pairwise comparisons of homogeneous and heterogenized forests on γ-, α-, and β-level with focus on rare (q=0), common (q=1) and dominant species (q=2) as well as on taxonomic (TD), functional (FD) and phylogenetic diversity (PD) of 1308 beetle species in 234 patches. Positive values indicate higher diversity levels in ESBC districts compared to control districts. Error bars denote the 95% confidence intervals calculated from weighted averages of bootstrapped, district-specific diversity estimates for each of the 27 cases.

### β-deviation

To compute pairwise β-diversity indices (distance matrices) as a measure of dissimilarity for all study plots, we utilized the function iNEXTbeta3D_pair3D adapted from the iNEXT.beta3D package. This step was repeated for a randomized data set generated from the observed data as a reference null model. For each site, all observed individuals were shuffled between the patches of the site and thus randomly redistributed within the ESBC and control district of a site (Kraft *et al*. 2015; Mori *et al*. 2015b), preserving γ-diversity and the relative abundance of each species in a site. β-deviation was calculated as the difference of β-diversities between observed and randomized data. Increasing positive β-deviation indicates a greater difference between the higher observed dissimilarity and the lower expected dissimilarity of the null model, suggesting that patches differ more in their species composition than expected under random community assembly.

### Linear mixed model

To model the β-deviation for all nine facets of diversity (taxonomic, phylogenetic, and functional diversity in combination with Hill numbers q = 0, 1, and 2) we used the pairwise differences in forest structure, abiotic conditions, and spatial distance as fixed factors. We estimated the contribution of each predictor treatment specific (ESBC and control district). Using this in a single model allows a direct comparison of the effect sizes for each treatment. For testing for differences in the slopes we also present the full interaction model in the Appendix (Tab. S1). The final models were fitted using the function *lmer* of the ‘lme4’ package (Bates et al. 2015). Rather than assuming that pairwise distances are independent observations (as in conventional fixed-effects models), we employed site as a random-effect to capture both within-district and among-district dependence (Harrison *et al*. 2018; Chao *et al*. 2024). We reported z-values as a standardized metrics of effect sizes.

To additionally analyse when functional diversity patterns reveal metacommunity assembly processes, we examined functional β-diversity independently from the changes in taxonomic diversity by using another null model approach. For 100 replicates of the null model, we permuted species labels across the species trait matrix, while the species abundance matrix and thus its spatial pattern was kept unchanged (Jeliazkov & Chase 2024). We calculated the standardized effect size (SES) normalizing observed functional β-diversity by mean and standard deviation of the randomized samples. Positive SES were considered to indicate higher functional turnover relative to taxonomic turnover, whereas a SES lower than 0 indicated lower functional turnover. Absolute differences greater than 1.96 indicate significant effects. Finally, we compared SES levels in control and ESBC districts by a GLMM again with site as a random factor.

## Results

### Diversity

Our final data set comprised 1308 beetle species (37 -143 per patch) in 103,714 specimens (153 -1888 per patch). The meta-analyses showed the most pronounced effect size in effective species number for γ-diversity in ESBC districts by an increase of about 60 species compared to control districts when focusing on rare species (q=0, Fig. 3). This was driven by both an increase in α-diversity of about 30 species and of β-diversity of about 10%. The effective number of common species (q=1) also increased at the landscape scale, but by 6 species only, again driven by α- and β-diversity. For dominant species (q=2), we found no effect on γ-diversity, but there was a small increase in α-diversity (0.8 species). Except for q=2 β-diversity of functional and phylogenetic diversities (effective numbers of functional groups and lineages) showed lower responses to ESBC forests than taxonomic diversity, which is an indication of functional and phylogenetic redundancy. Here the effects were most pronounced for β-diversity. From rare to dominant species, the increase in functional and phylogenetic β-diversity differences between ESBC and control districts turned into a significantly negative effect. Underlying district-specific results including local diversity estimates are presented in Supplement, Fig. S1-S27.

### β-deviation

Modelling of the β-deviation for all Hill numbers and facets of diversity showed strong positive effects of the manipulated differences between patches via forest structure in ESBC districts, for all Hill numbers of taxonomic diversity (Table 1). Space was significant here only for common species (q=1). The z-values of structure versus space were much larger in the ESBC districts, indicating the dominance of a *species-sorting* effect in the ESBC patches. In the control districts rare and common species were also affected by structural differences. However, in common and dominant species space was the most relevant predictor. Differences in abiotic conditions, using Ellenberg indicator values as a proxy for unmeasured spatial abiotic heterogeneity, did not significantly contribute to explaining taxonomic β-deviation, suggesting that most of the environmental heterogeneity was already captured by the experimentally manipulated and measured variables. Conditional R squared values were highest for taxonomic diversity (Tab. 1).

**Table 1:** Results of linear mixed interaction models of the β-deviation, i.e. the difference between β-diversity of observed vs. null-model data (with randomized distribution of individuals across patches in each site) for taxonomic (TD), functional (FD), and phylogenetic (PD) β-diversity in combination with Hill numbers q=0, 1, 2 (rare, common, and dominant species). As predictors we used the structural difference between patches (Structure), the difference in space (Spatial), and the difference in abiotic conditions (Abiotic) extracted from Ellenberg indicator values for both levels of deadwood and canopy structure (Control vs. ESBC), as well as Site as random factor. Last row provides the conditional R squared value of the model.

| Diversity facet | TD |  |  | FD |  |  | PD |  |  |
| --- | --- | --- | --- | --- | --- | --- | --- | --- | --- |
| Hill number q | 0 | 1 | 2 | 0 | 1 | 2 | 0 | 1 | 2 |
| Control:Structure | **3.628** | **3.196** | 1.579 | -0.068 | 0.352 | 0.340 | **2.002** | 0.791 | 0.644 |
| ESBC:Structure | **4.936** | **11.75** | **7.685** | **3.865** | 0.509 | 0.495 | **2.372** | 1.722 | -0.254 |
| Control:Abiotic | -0.256 | -1.155 | -0.467 | 1.258 | **-2.196** | -1.673 | -0.471 | **-2.369** | **-2.347** |
| ESBC:Abiotic | 1.124 | 1.152 | 1.807 | **-2.007** | 0.530 | 0.790 | **2.373** | 1.476 | 1.040 |
| Control:Spatial | 0.835 | **6.994** | **7.229** | 0.878 | **5.844** | **6.287** | 0.889 | **6.909** | **6.987** |
| ESBC:Spatial | 1.538 | **3.092** | 1.715 | 0.017 | 1.230 | 1.340 | **2.345** | 1.313 | 1.220 |
| R2 conditional | 0.11 | 0.35 | 0.38 | 0.071 | 0.15 | 0.18 | 0.10 | 0.17 | 0.17 |

The β-deviation of functional and phylogenetic diversity revealed similar patterns. In the treatment districts the structural heterogeneity increased functional/phylogenetic diversity of rare species, additionally supporting the *species-sorting* effect due to differences in species’ attributes. In homogeneous control districts, however, functional and phylogenetic diversity of common and dominant species increased with increasing spatial distance, which supports the *patch-dynamics* archetype with species showing differences in dispersal and competition. Phylogenetic β-deviation in heterogeneous forests was the only one which increased for rare species with difference in abiotic conditions between patches, which supports the view that the gain in phylogenetic diversity for rare species (Fig. 3) is promoted not only by pure light or deadwood as quantified for the structure, but also for more subtle gradients, e.g. of moisture. Standardized effect sizes of mean pairwise distances in functional β-diversity significantly decreased from control districts to ESBC districts for common and rare species (Supplement, Table S3) indicating a transition towards metacommunities more affected by habitat filtering even when controlling for taxonomic diversity.

## Discussion

Our results supported our first hypothesis that heterogenization of a homogeneous temperate production forest increases the diversity of beetles but only with focus on rare and common species, while dominant species remained unaffected or were even disfavoured. The finding that the increase in γ-diversity was determined by an increase in α- and β-diversity confirmed the predictions that both *mass-effects* and *species-sorting* contribute to the higher γ-diversity. This was also supported by the increase in functional or phylogenetic diversity again on the α-scale. Furthermore, the consistent result of larger effects of space on β-deviation in homogeneous forests shifting to reduced spatial effects, but an increase of forest structure effects in heterogeneous forests, supported that ESBC forests are more driven by *species-sorting* processes rather than *patch-dynamics*. This could particularly be observed for functional and phylogenetic diversity which reflect species differences in habitat requirements and mobility and thus are linked to *species-sorting*.

Our study implements the concluding suggestion of Jeliazkov and Chase (2024) to consider scale-explicit approaches depending on disturbance, scales, and decomposition into α-, γ-, and specifically β-components for improving metacommunity analysis. However, a particular problem in β-diversity analyses has been observed for many years. In observational data a fraction of β-response can be regularly attributed to locally varying unobserved species resulting in artifacts of the analysis which distort the results (Tuomisto 2010). This issue is even more pronounced in experimental approaches with substantial, independent variation in predictors where sample coverage has been shown to systematically correlate with treatment (Kortmann *et al*. 2025b; Rothacher *et al*. 2025) which is increasingly relevant in highly diverse taxa with many unobserved species, as in beetles. It does not only matter in species-rich and structurally extremely heterogeneous tropical forests, but also in European temperate zones. Recent analyses showed that many studies with insect samples from temperate zones are incomplete, in cases using flight and pitfall traps in beetles, light trapping of moths or even for total insect data from malaise traps (Müller & Brandl 2009; Kortmann *et al*. 2025a; Püls *et al*. 2025). In addition, it especially affects the conclusions from null model approaches which are evaluating the relative role of different assembly processes in a spatially hierarchical setting (Mori *et al*. 2015a). Tucker et al. (2016) demonstrated this effect of a higher β-diversity in undersampled communities with simulated data and suggested developing methods to control for it. This has been solved recently by standardizing not only diversity by the comprehensive approach of attribute diversity, but also dissimilarity between communities, by controlling for sample coverage (Chao *et al*. 2023). In contrast to the widely used Bray-Curtis-dissimilarity the generalized concept is applicable even to dissimilarities along Hill numbers and enables a much wider range of β-diversity applications. Moreover, this unified approach for taxonomic, functional, and phylogenetic diversity, does not only allow direct comparisons of the different units, but also to generalize the β-diversity concept to the different aspects of species dissimilarity as previously proposed by Logue *et al*. (2011). Finally, the difference is not only reflected in traits. Besides species identity, species abundance substantially affects population dynamics and community assembly. Larger populations of dominant species have, for instance, higher dispersal potential than smaller populations and are less affected in survival by negative stochastic effects even if sometimes most affected in general (see for insects in Müller *et al*. 2024; van Klink *et al*. 2024). The use of Hill numbers in our study provides a generalized framework to investigate metacommunity patterns from different angles without any additional information about species attributes (see also Guzman *et al*. 2022).

### Patch-dynamics archetype

The homogeneous forests are mostly in accordance with metacommunities dominated by *patch dynamics* (Levins & Culver 1971; Leibold *et al*. 2004). By centuries of human management, they are rather homogeneous in terms of tree dimension, vertical and horizontal structure. The patches within such a managed forest are rather similar, and we can expect community differences mainly to be explained by distance (Holyoak *et al*. 2005). Metacommunities in homogeneous environments are determined by differences in species ability to disperse, to colonize and to outcompete other species. Beetles show a wide range in the dispersal ability from flightless species to species dispersing actively over many kilometres (Buse 2012; Komonen & Müller 2018). Even though the maximum distance between two patches within a district was only ∼1km, which might be bridged by all species in our data set (Komonen & Müller 2018), we found a clear spatial effect independent from environmental differences in both homogeneous and heterogeneous forests. This was supported even for the direct measures of functional and phylogenetic β-deviation. *Patch dynamics* occurring in both forest types but being more pronounced in homogeneous forests suggests that this mechanism always contributes to metacommunities of beetles in forests but dominates in homogeneous ones. Even if we cannot compare directly, β-deviation analyses of tree species in continuous tropical old-growth forests also revealed a predominant effect of space (Myers *et al*. 2013).

The beetles in our data do not only differ in their abundance and mobility (from flightless to high mobile), but also in their ability to outcompete other species after colonization. One prominent example is Ambrosia beetles. They have evolved highly flexible life history strategies during the colonization of new patches, e.g. by sib-mating without inbreeding depression, changing the level of social behaviour, and adopting a complex mutualism with their ambrosia fungi (Biedermann 2010; Biedermann *et al*. 2011). In contrast, other beetle species are less flexible in traits resulting in a gradient of species’ location on the trade-off association between dispersal ability and competitive strength, which is required for *patch-dynamics* (Logue *et al*. 2011).

### Species-sorting

MacArthur and MacArthur (1961) laid the cornerstone for the habitat heterogeneity hypothesis using the gradient of vertical heterogeneity of forests as an axis of different niches for bird species. Later this single gradient of habitat heterogeneity has been expanded to a number of biotic and abiotic heterogeneity dimensions in forests with horizontal heterogeneity as one of the overall most prominent ones among taxa and functions (Heidrich *et al*. 2020). In this light, our substantial manipulation of forest structure, including the horizontal heterogeneity and amount of deadwood should end up in species-sorting (Leibold *et al*. 2004; Van der Gucht *et al*. 2007; Hernandez-Ordonez *et al*. 2019). We expected this especially, because we manipulated deadwood, which promotes many obligatory, but also facultative saproxylic arthropod and particularly beetle species (Seibold *et al*. 2016; Graf *et al*. 2022). In fact, the increase in β-diversity and β-deviation with increasing forest structure differences, supports this at least for rare and common species. The question remains, why we could not find an effect on diversity of dominant species. Here we must note that our local interventions did not change the structure of deadwood and light availability of large, forested landscapes. So, the majority of the forest remained even-aged, closed-canopy, and homogeneous making specific species the dominant species with high abundancies resistant to the ESBC treatment. However, dominant species can be affected by habitat heterogeneity under specific conditions. An analysis of the rarity level of the same forest beetles in Europe in response to wilderness areas revealed that rare species in Europe were dominant in the wilderness areas of Mongolia and vice versa (Müller *et al*. 2013). Similarly, we have reports of single species, e.g., the highly endangered *Peltis grossa* (Trogossitidae) formerly very rare due to modern forestry over centuries, turning into a dominant species after landscape-wide pulses of deadwood in large open forests after windstorm and bark beetle disturbances (Busse *et al*. 2022). Together with our findings, this shows that our patch-wide interventions were insufficient to affect the *species-sorting* of dominant species in production forests of the temperate European zone.

### Mass-effect

The second archetype commonly stressed in metacommunity theory with increasing heterogeneity is the *mass-effect* stimulating source-sink dynamics (Mouquet & Loreau 2003; Leibold *et al*. 2004). It has been stated that without information of dispersal rates or frequencies it cannot be distinguished from the *species-sorting* effect (Logue *et al*. 2011). However, our comprehensive approach on different spatial scales provides some support that the *mass-effect* is a critical mechanism providing higher γ-diversity in ESBC forests. We found a clear effect of the treatment on α-diversity. This indicates that the gain of the resources deadwood and sunlight in the former homogeneous, dark forest locally increases the diversity of beetles. Considering that spatial distance additionally affected the β-deviation beyond the pronounced effects of forest structure suggests that at least the increase in common species is also triggered by *mass-effects*. In addition, it could be responsible for the slightly negative effect in β-diversity in dominant species (Fig. 3), when massive dispersal of the most frequent species is promoted eventually resulting in homogenization of species distribution (Mouquet & Loreau 2002; Pardini *et al*. 2010). We could not identify such a spatial effect on β-deviation for rare species, but we found a strong effect on α-diversity of rare species. This might be explained by the fact that many rare species are promoted by resource pulses at the α-scale, but as they are too rare, this does not display in β-deviations. Nevertheless, the large contribution of α-diversity to higher γ-diversity in ESBC districts cannot be explained only by β-diversity suggesting an overall *mass-effect*.

### Facets of diversity

For maximum significance of functional diversity Jeliazkov and Chase (2024) suggested the selection of traits related to reproduction and mobility. Thus, we combined body size and feeding strategy with species response to light (brightness level) and to deadwood (saproxylic guild) which have been proven to be sensitive for environmental gradients as manipulated in our experiment (e.g. Gossner *et al*. 2013). Like Jeliazkov & Chase (2024), we found that in general diversity patterns were explained best for taxonomic diversity. We share their sceptical perspective on the potential explanatory contribution of functional diversity in addition to taxonomic diversity for rare species as SES do not indicate trait divergence or convergence when controlled for taxonomic diversity. However, for common and dominant species mostly positive SES values hint on trait divergence. In contrast to Jeliazkov and Chase (2024) we based our SES calculation on the multiplicative Hill number approach for diversity decomposition (Chao *et al*. 2023) which allows to differentiate the perspective between rare and common species in addition to dominant species. When considering not only the total explained variance of a model, but also the relevance of individual predictors in comparison to each other, nuanced insights can be gained from the differences between PD and FD as well as between the Hill numbers. For FD and PD forest structure affects mainly rare species indicating *species-sorting*, while the impact of space prevails in common and dominant species where successful dispersal can be triggered by abundance and constitute a *mass-effect*. Despite the general correlation between effect sizes in PD and FD, there are also deviating cases, e.g., in rare species and the impact of space in ESBC habitat, eventually indicating opposing effects of different traits aggregated in functional diversity.

### From mechanisms to management strategies

At the midpoint of the UN decade of restoration, success is still limited. From a forest management perspective, increasingly frequent natural disturbances are used as guidelines for new strategies aiming to improve resistance and biodiversity of forests (Aszalós *et al*. 2022). Many of these strategies aim to locally increase the number of old-growth attributes, such as trees with microhabitats or deadwood (Bauhus *et al*. 2009) or to mimic natural disturbance events (Seibold *et al*. 2016). However, natural events provide no standardized design, are most often not replicated, and spatially not independent. These factors prevent the necessary comprehensive understanding of the effects on diversity and the essential mechanisms of metacommunity theory. In addition, they increase the risk of misinterpretation and might even contribute to establish unproven paradigms (Weisser *et al*. 2023). In contrast to long lasting expectations two large global assessments recently found that land use intensity does not consistently form homogeneous communities, and that higher γ-diversity is often driven by increasing α-diversity, while effects of β-diversity were less consistent (Gonçalves-Souza & Sanders 2025; Keck *et al*. 2025). While the statistical tools to identify mechanisms along the axes of different species, habitat difference, and dispersal seem to be available now, more replicated studies in which homogeneous landscapes are experimentally heterogenized are necessary to decouple potential drivers and improve the understanding of decline or restoration success particularly for invertebrates

### Outlook

Taking beetles as a highly diverse model group, we derive several important implications for restoration of homogeneous temperate production forests. First, canopy openings by gap felling, group wise dieback of trees by insects, summer storms, or droughts and the enhancement of deadwood do not affect the diversity of dominant beetles in temperate forests. Second, all local resource pulses by light and deadwood increase not only the α-diversity but also translates into a higher γ-diversity of rare and common beetle species, probably by *mass-effects*. Additionally, increasing the between-patch heterogeneity also increases γ-diversity of rare and common species most probably by *species-sorting* effects. Therefore, both strategies, an opportunistic increase of resources and a targeted spatial heterogeneity in forests seem very promising in restoring the beetle diversity of homogeneous production forests, which could be extended to other taxa.

## Acknowledgements and funding

We thank Elisa Stengel, Luisa Pflumm, Thorben Riehe, Michael Junginger, Boris Büche, Alexander Szallies, Ruth Pickert, Sophia Hochrein, Katharina Kallnik, Marina Wolz, Manuel Mauermann, Bernd Anell, Jens Geigenmüller, Hans Bahr, Interns of the Bavarian Forest Nationalpark and all other assistants in the field and laboratory. We also thank the entire BETA-FOR team for their support. This work was supported by the Deutsche Forschungsgemeinschaft [BETA-FOR, 459717468]. J.R. received funding from the Bavarian State Ministry for Food, Agriculture, Forestry and Tourism (grant no. L062).

