## Supplement for "Old growth attributes by chain saw: how between-patch heterogeneity changes the metacommunities of beetles in temperate forests"

Fig. S1 to S27: All, i.e. in total 27, meta-analyses of taxonomic (TD), phylogenetic (PD), and functional (FD) diversity (3x) at coverage level of 0.95 for Hill numbers q=0, 1, and 2 (3x) at alpha, gamma, and beta (1-S) scale (3x), each for eleven study sites. The bottom row of each figure indicates the aggregated results at the meta-analysis level with bootstrapping confidence levels (lower LCL, upper UCL) indicating overall significance when not including 0. W(fixed) in each row indicates weighting of each site within the meta-analysis according to site-specific confidence intervals. Column Difference gives the difference between estimated diversity for Enhanced ESBC and Control districts. These analyses are summarized in Fig. 3 of the main manuscript.

**Alpha with Coverage = 0.95**

| Site | TD(q=0) Enhanced | TD(q=0) Control | Difference | LCL | UCL | W(fixed) |
|---|---|---|---|---|---|---|
| B04 | 127.721 | 80.161 | 47.561 | 33.47 | 61.652 | 9.1% |
| B05 | 134.019 | 128.59 | 5.429 | −10.189 | 21.047 | 7.41% |
| B06 | 98.816 | 89.523 | 9.293 | −4.773 | 23.358 | 9.14% |
| B07 | 108.028 | 55.51 | 52.518 | 42.434 | 62.602 | 17.77% |
| H09 | 109.694 | 76.586 | 33.109 | 16.014 | 50.203 | 6.18% |
| L11 | 127.894 | 80.684 | 47.21 | 29.393 | 65.027 | 5.69% |
| P08 | 150.04 | 129.981 | 20.059 | 3.615 | 36.503 | 6.68% |
| S10 | 100.687 | 81.678 | 19.01 | 7.239 | 30.78 | 13.05% |
| U01 | 173.954 | 133.012 | 40.942 | 27.209 | 54.675 | 9.58% |
| U02 | 178.462 | 134.919 | 43.543 | 28.511 | 58.575 | 8% |
| U03 | 182.073 | 157.026 | 25.047 | 9.413 | 40.682 | 7.39% |
| **Meta analysis** | | | **32.728** | **28.477** | **36.979** | |

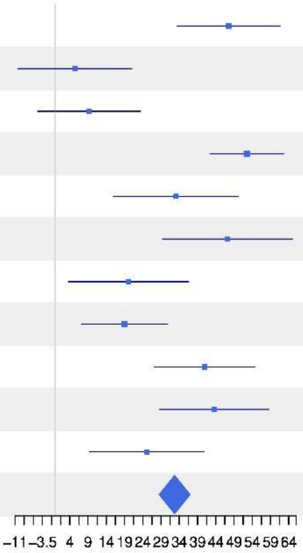


**Alpha with Coverage = 0.95**

| Site | TD(q=1) Enhanced | TD(q=1) Control | Difference | LCL | UCL | W(fixed) |
|---|---|---|---|---|---|---|
| B04 | 20.742 | 16.296 | 4.445 | 2.868 | 6.023 | 10.36% |
| B05 | 27.24 | 40.528 | −13.288 | −16.063 | −10.514 | 3.35% |
| B06 | 15.448 | 13.383 | 2.064 | 0.87 | 3.258 | 18.09% |
| B07 | 12.701 | 7.04 | 5.661 | 4.865 | 6.456 | 40.74% |
| H09 | 32.301 | 22.158 | 10.143 | 7.758 | 12.528 | 4.53% |
| L11 | 45.822 | 14.91 | 30.912 | 27.259 | 34.566 | 1.93% |
| P08 | 23.64 | 32.536 | −8.896 | −11.047 | −6.745 | 5.57% |
| S10 | 23.879 | 24.692 | −0.814 | −2.736 | 1.108 | 6.98% |
| U01 | 57.149 | 27.013 | 30.136 | 27.704 | 32.568 | 4.36% |
| U02 | 62.756 | 32.132 | 30.624 | 27.324 | 33.924 | 2.37% |
| U03 | 57.39 | 54.471 | 2.919 | −0.971 | 6.809 | 1.7% |
| **Meta analysis** | | | **5.289** | **4.781** | **5.797** | |

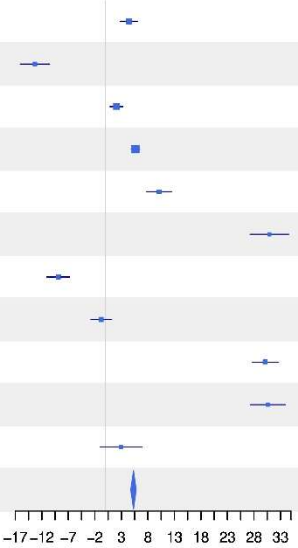

### Alpha with Coverage = 0.95

| Site | TD(q=2) Enhanced | TD(q=2) Control | Difference | LCL | UCL | W(fixed) |
|---|---|---|---|---|---|---|
| B04 | 6.482 | 6.849 | −0.367 | −0.948 | 0.213 | 7.71% |
| B05 | 8.27 | 17.295 | −9.024 | −10.311 | −7.738 | 1.57% |
| B06 | 5.961 | 4.617 | 1.344 | 0.97 | 1.719 | 18.55% |
| B07 | 3.736 | 2.65 | 1.086 | 0.877 | 1.294 | 59.74% |
| H09 | 17.024 | 10.137 | 6.888 | 5.667 | 8.108 | 1.75% |
| L11 | 18.55 | 5.372 | 13.178 | 11.425 | 14.93 | 0.85% |
| P08 | 2.624 | 13.224 | −10.6 | −11.405 | −9.795 | 4.01% |
| S10 | 8.81 | 10.941 | −2.131 | −3.092 | −1.17 | 2.81% |
| U01 | 20.781 | 6.026 | 14.754 | 13.441 | 16.067 | 1.51% |
| U02 | 22.543 | 7.346 | 15.197 | 13.398 | 16.996 | 0.8% |
| U03 | 18.735 | 20.741 | −2.006 | −3.937 | −0.075 | 0.7% |
| **Meta analysis** | | | **0.805** | **0.644** | **0.967** | |

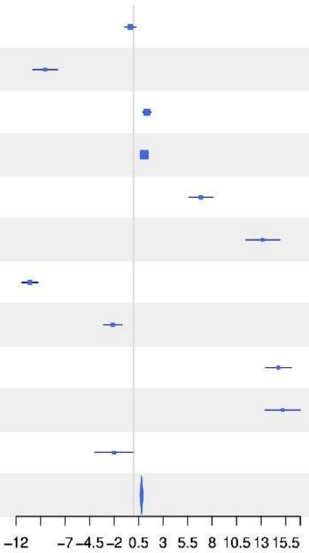


### Gamma with Coverage = 0.95

| Site | TD(q=0) Enhanced | TD(q=0) Control | Difference | LCL | UCL | W(fixed) |
|---|---|---|---|---|---|---|
| B04 | 177.424 | 102.738 | 74.686 | 54.8 | 94.572 | 10.88% |
| B05 | 210.24 | 198.799 | 11.442 | −12.377 | 35.261 | 7.58% |
| B06 | 137.771 | 106.836 | 30.934 | 14.733 | 47.136 | 16.39% |
| B07 | 148.443 | 70.203 | 78.24 | 64.712 | 91.769 | 23.51% |
| H09 | 169.124 | 112.93 | 56.194 | 32.583 | 79.805 | 7.72% |
| L11 | 267.236 | 107.256 | 159.98 | 124.816 | 195.143 | 3.48% |
| P08 | 215.451 | 168.128 | 47.323 | 25.712 | 68.934 | 9.21% |
| S10 | 146.474 | 125.804 | 20.67 | −0.504 | 41.844 | 9.6% |
| U01 | 360.988 | 256.097 | 104.891 | 73.085 | 136.697 | 4.25% |
| U02 | 425.328 | 242.372 | 182.956 | 149.286 | 216.625 | 3.8% |
| U03 | 425.248 | 275.908 | 149.34 | 114.717 | 183.962 | 3.59% |
| **Meta analysis** | | | **65.464** | **58.905** | **72.023** | |

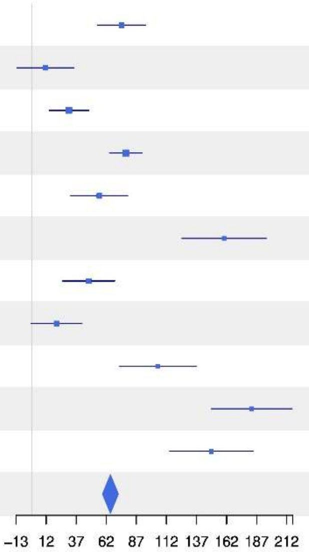

### Gamma with Coverage = 0.95

| Site | TD(q=1) Enhanced | TD(q=1) Control | Difference | LCL | UCL | W(fixed) |
|---|---|---|---|---|---|---|
| B04 | 25.384 | 19.804 | 5.58 | 3.231 | 7.929 | 8.41% |
| B05 | 34.144 | 53.106 | −18.962 | −22.533 | −15.391 | 3.64% |
| B06 | 21.054 | 14.962 | 6.092 | 4.669 | 7.515 | 22.93% |
| B07 | 15.537 | 10.686 | 4.851 | 3.831 | 5.871 | 44.63% |
| H09 | 39.811 | 28.555 | 11.256 | 8.368 | 14.144 | 5.57% |
| L11 | 69.047 | 17.462 | 51.584 | 46.08 | 57.089 | 1.53% |
| P08 | 45.067 | 42.413 | 2.654 | −0.714 | 6.021 | 4.09% |
| S10 | 29.865 | 32.599 | −2.734 | −5.726 | 0.258 | 5.19% |
| U01 | 99.511 | 52.324 | 47.186 | 41.99 | 52.383 | 1.72% |
| U02 | 119.887 | 56.095 | 63.793 | 57.585 | 70 | 1.2% |
| U03 | 110.448 | 83.254 | 27.194 | 20.637 | 33.751 | 1.08% |
| **Meta analysis** | | | **6.599** | **5.917** | **7.28** | |

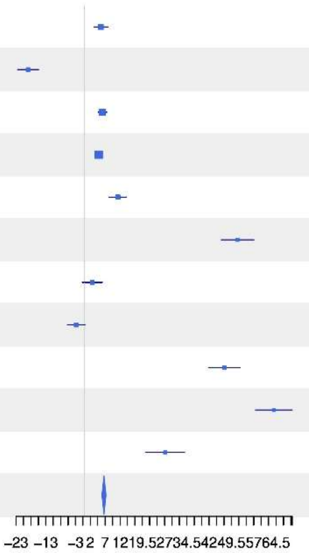


### Gamma with Coverage = 0.95

| Site | TD(q=2) Enhanced | TD(q=2) Control | Difference | LCL | UCL | W(fixed) |
|---|---|---|---|---|---|---|
| B04 | 7.145 | 8.096 | −0.952 | −1.607 | −0.296 | 9.83% |
| B05 | 9.76 | 26.596 | −16.836 | −18.501 | −15.171 | 1.52% |
| B06 | 8.187 | 4.921 | 3.266 | 2.743 | 3.79 | 15.41% |
| B07 | 4.304 | 4.504 | −0.201 | −0.456 | 0.055 | 64.58% |
| H09 | 21.861 | 13.657 | 8.204 | 6.727 | 9.681 | 1.94% |
| L11 | 26.976 | 6.225 | 20.751 | 17.981 | 23.521 | 0.55% |
| P08 | 13.771 | 17.88 | −4.109 | −5.395 | −2.824 | 2.55% |
| S10 | 10.957 | 14.647 | −3.69 | −5.002 | −2.378 | 2.45% |
| U01 | 39.779 | 18.85 | 20.929 | 18.017 | 23.841 | 0.5% |
| U02 | 48.974 | 22.395 | 26.579 | 23.065 | 30.094 | 0.34% |
| U03 | 43.154 | 39.179 | 3.975 | 0.335 | 7.616 | 0.32% |
| **Meta analysis** | | | **0.309** | **0.104** | **0.515** | |

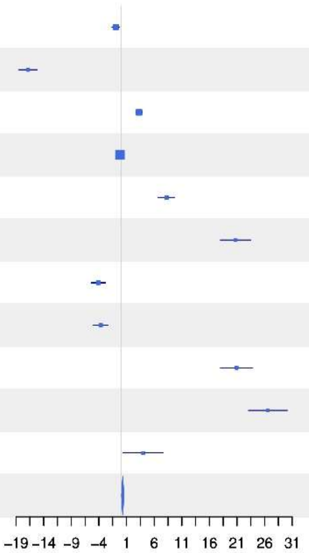

**1−S with Coverage = 0.95**

| Site | TD(q=0) Enhanced | TD(q=0) Control | Difference | LCL | UCL | W(fixed) |
|---|---|---|---|---|---|---|
| B04 | 0.315 | 0.247 | 0.068 | 0.025 | 0.111 | 5.68% |
| B05 | 0.408 | 0.397 | 0.011 | −0.025 | 0.046 | 8.54% |
| B06 | 0.318 | 0.182 | 0.136 | 0.086 | 0.185 | 4.3% |
| B07 | 0.306 | 0.235 | 0.071 | 0.016 | 0.126 | 3.45% |
| H09 | 0.395 | 0.362 | 0.033 | −0.018 | 0.085 | 4% |
| L11 | 0.587 | 0.279 | 0.308 | 0.266 | 0.349 | 6.14% |
| P08 | 0.342 | 0.255 | 0.086 | 0.048 | 0.124 | 7.35% |
| S10 | 0.352 | 0.395 | −0.043 | −0.094 | 0.008 | 4.08% |
| U01 | 0.555 | 0.515 | 0.04 | 0.017 | 0.064 | 19.04% |
| U02 | 0.622 | 0.475 | 0.147 | 0.123 | 0.171 | 18.81% |
| U03 | 0.613 | 0.462 | 0.151 | 0.127 | 0.175 | 18.61% |
| **Meta analysis** | | | **0.101** | **0.091** | **0.112** | |

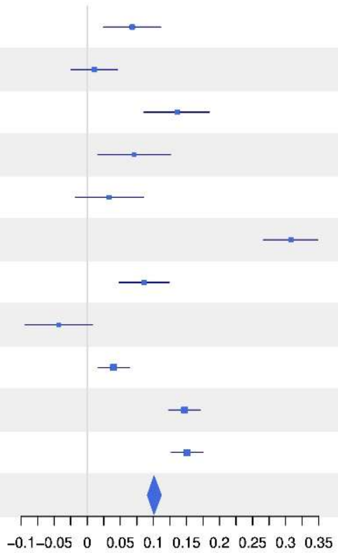


**1−S with Coverage = 0.95**

| Site | TD(q=1) Enhanced | TD(q=1) Control | Difference | LCL | UCL | W(fixed) |
|---|---|---|---|---|---|---|
| B04 | 0.206 | 0.199 | 0.006 | −0.026 | 0.039 | 5.4% |
| B05 | 0.227 | 0.266 | −0.039 | −0.069 | −0.009 | 6.5% |
| B06 | 0.3 | 0.119 | 0.181 | 0.152 | 0.21 | 6.83% |
| B07 | 0.205 | 0.384 | −0.178 | −0.201 | −0.156 | 11.1% |
| H09 | 0.212 | 0.252 | −0.04 | −0.07 | −0.009 | 6.27% |
| L11 | 0.378 | 0.164 | 0.214 | 0.176 | 0.252 | 4.01% |
| P08 | 0.535 | 0.262 | 0.273 | 0.247 | 0.299 | 8.81% |
| S10 | 0.225 | 0.273 | −0.047 | −0.08 | −0.015 | 5.45% |
| U01 | 0.456 | 0.518 | −0.062 | −0.081 | −0.044 | 17.28% |
| U02 | 0.511 | 0.458 | 0.053 | 0.034 | 0.072 | 15.94% |
| U03 | 0.515 | 0.37 | 0.144 | 0.123 | 0.166 | 12.42% |
| **Meta analysis** | | | **0.033** | **0.026** | **0.041** | |

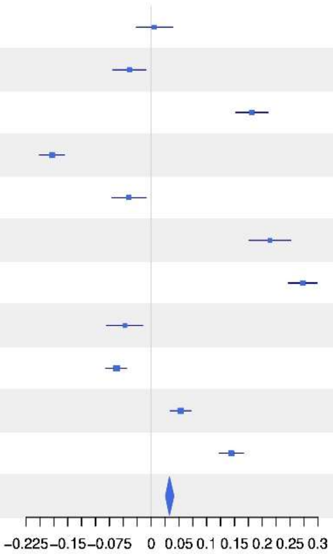

#### 1−S with Coverage = 0.95

| Site | TD(q=2) Enhanced | TD(q=2) Control | Difference | LCL | UCL | W(fixed) |
|---|---|---|---|---|---|---|
| B04 | 0.104 | 0.173 | −0.069 | −0.095 | −0.043 | 13.82% |
| B05 | 0.172 | 0.393 | −0.222 | −0.264 | −0.179 | 5.25% |
| B06 | 0.306 | 0.07 | 0.236 | 0.208 | 0.265 | 11.45% |
| B07 | 0.148 | 0.463 | −0.315 | −0.344 | −0.286 | 11.25% |
| H09 | 0.249 | 0.29 | −0.041 | −0.084 | 0.002 | 5.06% |
| L11 | 0.351 | 0.154 | 0.197 | 0.139 | 0.255 | 2.81% |
| P08 | 0.911 | 0.293 | 0.618 | 0.588 | 0.648 | 10.67% |
| S10 | 0.22 | 0.285 | −0.064 | −0.106 | −0.022 | 5.36% |
| U01 | 0.512 | 0.729 | −0.217 | −0.244 | −0.19 | 13.1% |
| U02 | 0.578 | 0.72 | −0.142 | −0.169 | −0.115 | 12.95% |
| U03 | 0.606 | 0.504 | 0.102 | 0.068 | 0.136 | 8.28% |
| **Meta analysis** | | | **−0.002** | **−0.012** | **0.008** | |

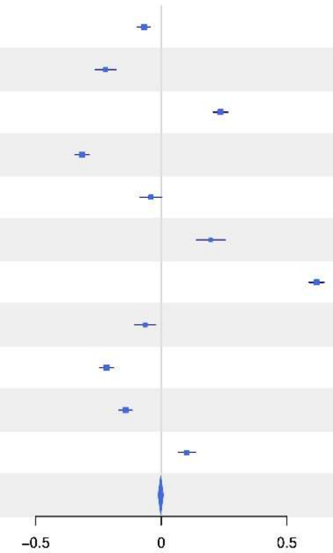


#### Alpha with Coverage = 0.95

| Site | PD(q=0) Enhanced | PD(q=0) Control | Difference | LCL | UCL | W(fixed) |
|---|---|---|---|---|---|---|
| B04 | 12.951 | 11.367 | 1.585 | 0.641 | 2.528 | 8.29% |
| B05 | 13.126 | 13.279 | −0.153 | −1.113 | 0.807 | 8.01% |
| B06 | 12.041 | 11.607 | 0.434 | −0.422 | 1.29 | 10.08% |
| B07 | 12.603 | 9.806 | 2.797 | 2.014 | 3.579 | 12.06% |
| H09 | 11.841 | 10.43 | 1.411 | 0.404 | 2.417 | 7.29% |
| L11 | 13.37 | 11.564 | 1.806 | 0.517 | 3.095 | 4.45% |
| P08 | 13.535 | 13.363 | 0.172 | −0.652 | 0.996 | 10.87% |
| S10 | 11.405 | 11.251 | 0.154 | −0.694 | 1.002 | 10.26% |
| U01 | 13.718 | 13.854 | −0.136 | −0.922 | 0.65 | 11.95% |
| U02 | 13.493 | 12.902 | 0.59 | −0.359 | 1.54 | 8.19% |
| U03 | 14.33 | 13.555 | 0.775 | −0.155 | 1.705 | 8.54% |
| **Meta analysis** | | | **0.816** | **0.545** | **1.088** | |

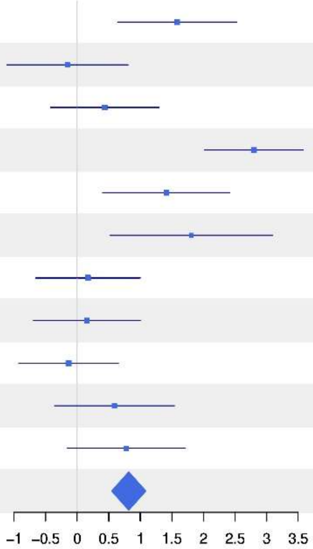

#### Alpha with Coverage = 0.95

| Site | PD(q=1) Enhanced | PD(q=1) Control | Difference | LCL | UCL | W(fixed) |
|---|---|---|---|---|---|---|
| B04 | 4.44 | 4.454 | −0.015 | −0.157 | 0.127 | 7.21% |
| B05 | 4.933 | 5.541 | −0.608 | −0.747 | −0.469 | 7.52% |
| B06 | 4.115 | 4.059 | 0.057 | −0.065 | 0.179 | 9.77% |
| B07 | 3.826 | 2.951 | 0.875 | 0.762 | 0.988 | 11.31% |
| H09 | 4.972 | 4.759 | 0.213 | 0.084 | 0.342 | 8.67% |
| L11 | 5.045 | 4.565 | 0.48 | 0.292 | 0.669 | 4.08% |
| P08 | 3.781 | 4.542 | −0.76 | −0.869 | −0.651 | 12.21% |
| S10 | 4.887 | 5.029 | −0.143 | −0.274 | −0.011 | 8.35% |
| U01 | 5.223 | 3.929 | 1.294 | 1.183 | 1.406 | 11.68% |
| U02 | 4.965 | 4.298 | 0.666 | 0.541 | 0.791 | 9.28% |
| U03 | 5.189 | 5.144 | 0.045 | −0.076 | 0.166 | 9.92% |
| **Meta analysis** | | | **0.209** | **0.17** | **0.247** | |

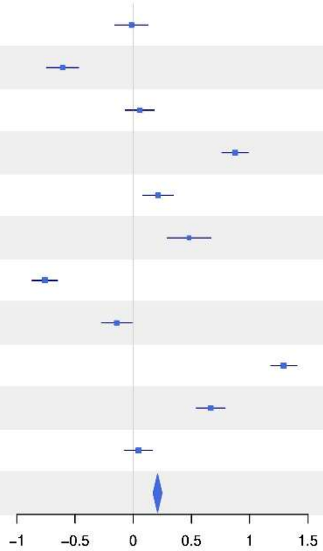


#### Alpha with Coverage = 0.95

| Site | PD(q=2) Enhanced | PD(q=2) Control | Difference | LCL | UCL | W(fixed) |
|---|---|---|---|---|---|---|
| B04 | 2.992 | 3.15 | −0.158 | −0.244 | −0.072 | 11.14% |
| B05 | 3.296 | 3.831 | −0.535 | −0.65 | −0.42 | 6.26% |
| B06 | 2.893 | 2.771 | 0.121 | 0.03 | 0.213 | 9.92% |
| B07 | 2.506 | 1.666 | 0.84 | 0.77 | 0.911 | 16.83% |
| H09 | 3.715 | 3.479 | 0.237 | 0.121 | 0.352 | 6.26% |
| L11 | 3.406 | 3.17 | 0.237 | 0.093 | 0.38 | 4.04% |
| P08 | 1.591 | 2.922 | −1.331 | −1.427 | −1.235 | 9.09% |
| S10 | 3.562 | 3.732 | −0.169 | −0.285 | −0.054 | 6.25% |
| U01 | 3.44 | 2.142 | 1.298 | 1.218 | 1.378 | 13.02% |
| U02 | 3.065 | 2.563 | 0.502 | 0.407 | 0.597 | 9.15% |
| U03 | 3.316 | 3.214 | 0.102 | 0.001 | 0.204 | 8.04% |
| **Meta analysis** | | | **0.219** | **0.19** | **0.247** | |

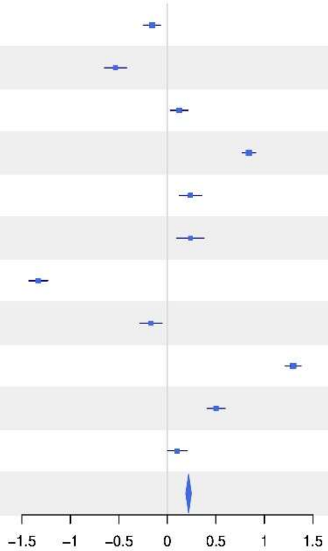

**Gamma with Coverage = 0.95**

| Site | PD(q=0) Enhanced | PD(q=0) Control | Difference | LCL | UCL | W(fixed) |
|---|---|---|---|---|---|---|
| B04 | 14.617 | 12.229 | 2.388 | 0.583 | 4.193 | 8.66% |
| B05 | 15.306 | 14.615 | 0.69 | −1.203 | 2.584 | 7.87% |
| B06 | 13.338 | 12.191 | 1.148 | −0.587 | 2.882 | 9.38% |
| B07 | 14.074 | 10.664 | 3.409 | 1.959 | 4.86 | 13.41% |
| H09 | 14.601 | 12.481 | 2.12 | −0.132 | 4.372 | 5.56% |
| L11 | 17.282 | 12.441 | 4.841 | 2.106 | 7.576 | 3.77% |
| P08 | 15.607 | 14.526 | 1.082 | −0.461 | 2.624 | 11.85% |
| S10 | 13.204 | 12.879 | 0.325 | −1.574 | 2.225 | 7.82% |
| U01 | 18.266 | 17.164 | 1.102 | −0.427 | 2.631 | 12.07% |
| U02 | 19.181 | 16.293 | 2.889 | 1.268 | 4.509 | 10.75% |
| U03 | 19.094 | 16.691 | 2.403 | 0.62 | 4.187 | 8.87% |
| **Meta analysis** | | | **1.937** | **1.405** | **2.468** | |

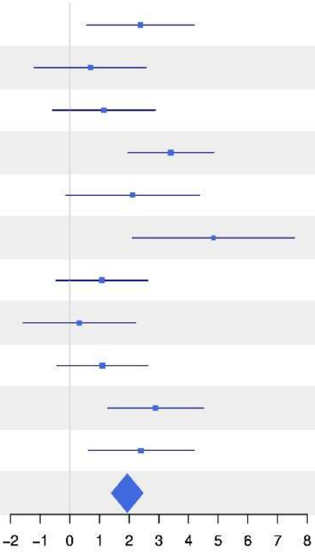


**Gamma with Coverage = 0.95**

| Site | PD(q=1) Enhanced | PD(q=1) Control | Difference | LCL | UCL | W(fixed) |
|---|---|---|---|---|---|---|
| B04 | 4.733 | 4.827 | −0.094 | −0.238 | 0.05 | 6.29% |
| B05 | 5.266 | 5.982 | −0.715 | −0.845 | −0.586 | 7.77% |
| B06 | 4.675 | 4.201 | 0.474 | 0.342 | 0.605 | 7.52% |
| B07 | 4.117 | 4.014 | 0.103 | −0.016 | 0.222 | 9.1% |
| H09 | 5.317 | 5.271 | 0.047 | −0.075 | 0.168 | 8.85% |
| L11 | 5.577 | 4.89 | 0.687 | 0.502 | 0.872 | 3.78% |
| P08 | 5.405 | 4.847 | 0.558 | 0.445 | 0.671 | 10.23% |
| S10 | 5.283 | 5.484 | −0.201 | −0.317 | −0.086 | 9.7% |
| U01 | 5.852 | 5.23 | 0.622 | 0.52 | 0.723 | 12.5% |
| U02 | 5.879 | 5.55 | 0.329 | 0.225 | 0.433 | 12.02% |
| U03 | 5.86 | 5.771 | 0.089 | −0.014 | 0.192 | 12.22% |
| **Meta analysis** | | | **0.179** | **0.143** | **0.215** | |

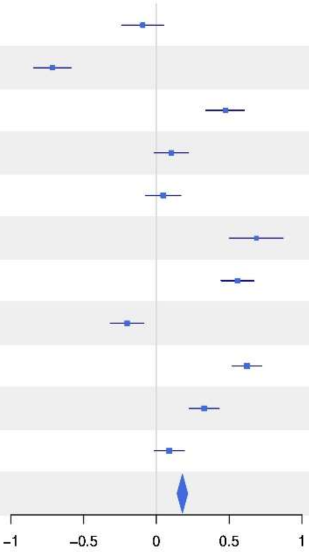

**Gamma with Coverage = 0.95**

| Site | PD(q=2) Enhanced | PD(q=2) Control | Difference | LCL | UCL | W(fixed) |
|---|---|---|---|---|---|---|
| B04 | 3.224 | 3.424 | −0.2 | −0.289 | −0.112 | 9.24% |
| B05 | 3.623 | 4.295 | −0.672 | −0.77 | −0.574 | 7.51% |
| B06 | 3.318 | 2.901 | 0.417 | 0.328 | 0.506 | 9.13% |
| B07 | 2.738 | 2.874 | −0.136 | −0.223 | −0.049 | 9.54% |
| H09 | 3.982 | 3.991 | −0.008 | −0.102 | 0.085 | 8.22% |
| L11 | 3.767 | 3.533 | 0.234 | 0.088 | 0.379 | 3.41% |
| P08 | 3.656 | 3.22 | 0.436 | 0.334 | 0.538 | 6.93% |
| S10 | 3.955 | 4.243 | −0.288 | −0.372 | −0.205 | 10.42% |
| U01 | 3.894 | 3.67 | 0.225 | 0.149 | 0.3 | 12.78% |
| U02 | 3.83 | 4.046 | −0.217 | −0.286 | −0.148 | 15.13% |
| U03 | 3.839 | 3.884 | −0.044 | −0.141 | 0.053 | 7.7% |
| **Meta analysis** | | | **−0.044** | **−0.071** | **−0.017** | |

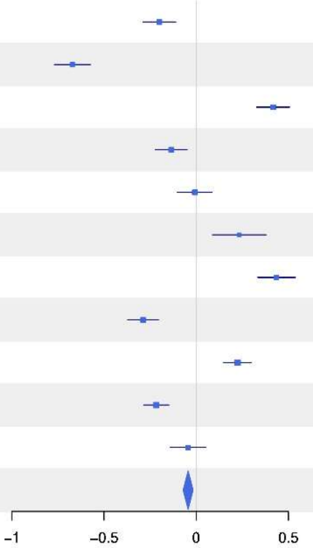


**1−S with Coverage = 0.95**

| Site | PD(q=0) Enhanced | PD(q=0) Control | Difference | LCL | UCL | W(fixed) |
|---|---|---|---|---|---|---|
| B04 | 0.128 | 0.079 | 0.049 | −0.014 | 0.112 | 7.73% |
| B05 | 0.16 | 0.103 | 0.057 | −0.002 | 0.117 | 8.59% |
| B06 | 0.109 | 0.054 | 0.056 | 0 | 0.112 | 9.82% |
| B07 | 0.118 | 0.091 | 0.027 | −0.03 | 0.084 | 9.53% |
| H09 | 0.213 | 0.185 | 0.028 | −0.048 | 0.104 | 5.3% |
| L11 | 0.255 | 0.079 | 0.175 | 0.087 | 0.264 | 3.93% |
| P08 | 0.149 | 0.09 | 0.059 | 0.003 | 0.116 | 9.66% |
| S10 | 0.153 | 0.142 | 0.011 | −0.05 | 0.072 | 8.19% |
| U01 | 0.267 | 0.207 | 0.06 | 0.014 | 0.107 | 14.15% |
| U02 | 0.318 | 0.223 | 0.095 | 0.04 | 0.15 | 10.28% |
| U03 | 0.267 | 0.201 | 0.066 | 0.017 | 0.115 | 12.83% |
| **Meta analysis** | | | **0.058** | **0.041** | **0.076** | |

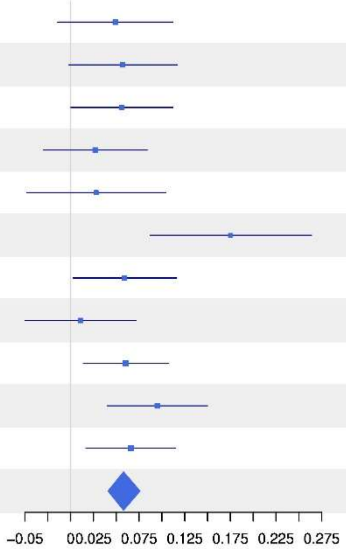

### 1−S with Coverage = 0.95

| Site | PD(q=1) Enhanced | PD(q=1) Control | Difference | LCL | UCL | W(fixed) |
|---|---|---|---|---|---|---|
| B04 | 0.07 | 0.087 | −0.017 | −0.031 | −0.003 | 9.48% |
| B05 | 0.071 | 0.083 | −0.012 | −0.025 | 0.002 | 9.71% |
| B06 | 0.135 | 0.038 | 0.096 | 0.084 | 0.109 | 11.19% |
| B07 | 0.079 | 0.298 | −0.218 | −0.231 | −0.205 | 10.53% |
| H09 | 0.073 | 0.109 | −0.036 | −0.052 | −0.02 | 6.89% |
| L11 | 0.107 | 0.075 | 0.032 | 0.013 | 0.052 | 4.77% |
| P08 | 0.338 | 0.071 | 0.267 | 0.255 | 0.279 | 12.94% |
| S10 | 0.084 | 0.093 | −0.009 | −0.026 | 0.008 | 5.87% |
| U01 | 0.115 | 0.267 | −0.151 | −0.165 | −0.138 | 10.47% |
| U02 | 0.167 | 0.242 | −0.075 | −0.09 | −0.06 | 7.64% |
| U03 | 0.123 | 0.116 | 0.006 | −0.007 | 0.019 | 10.51% |
| **Meta analysis** | | | **−0.003** | **−0.007** | **0.001** | |

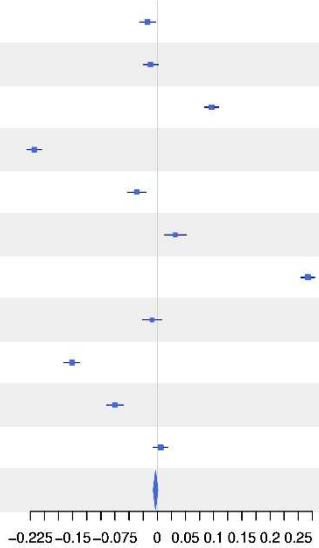


### 1−S with Coverage = 0.95

| Site | PD(q=2) Enhanced | PD(q=2) Control | Difference | LCL | UCL | W(fixed) |
|---|---|---|---|---|---|---|
| B04 | 0.081 | 0.09 | −0.009 | −0.024 | 0.006 | 14.34% |
| B05 | 0.101 | 0.122 | −0.02 | −0.041 | 0.001 | 7.61% |
| B06 | 0.144 | 0.05 | 0.094 | 0.077 | 0.111 | 11.77% |
| B07 | 0.095 | 0.473 | −0.378 | −0.397 | −0.359 | 8.89% |
| H09 | 0.075 | 0.144 | −0.069 | −0.093 | −0.045 | 5.71% |
| L11 | 0.108 | 0.116 | −0.008 | −0.032 | 0.016 | 5.84% |
| P08 | 0.635 | 0.104 | 0.531 | 0.515 | 0.548 | 11.78% |
| S10 | 0.112 | 0.136 | −0.024 | −0.047 | −0.001 | 6.01% |
| U01 | 0.125 | 0.446 | −0.321 | −0.339 | −0.303 | 10.23% |
| U02 | 0.214 | 0.393 | −0.179 | −0.2 | −0.158 | 7.45% |
| U03 | 0.146 | 0.185 | −0.039 | −0.056 | −0.021 | 10.38% |
| **Meta analysis** | | | **−0.019** | **−0.025** | **−0.013** | |

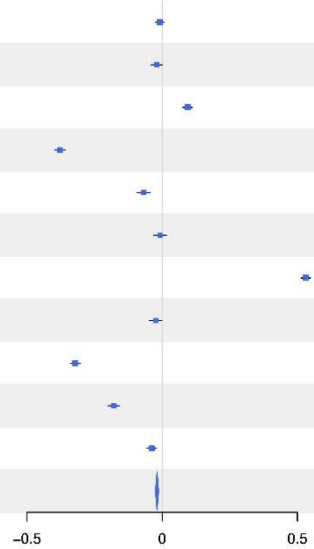

### Alpha with Coverage = 0.95

| Site | FD(q=0) Enhanced | FD(q=0) Control | Difference | LCL | UCL | W(fixed) |
|---|---|---|---|---|---|---|
| B04 | 11.079 | 9.525 | 1.554 | −0.186 | 3.294 | 7.08% |
| B05 | 11.688 | 11.474 | 0.214 | −1.272 | 1.7 | 9.71% |
| B06 | 9.736 | 9.578 | 0.158 | −1.16 | 1.476 | 12.33% |
| B07 | 10.623 | 7.989 | 2.634 | 1.174 | 4.094 | 10.06% |
| H09 | 10.675 | 9.129 | 1.546 | −0.082 | 3.174 | 8.09% |
| L11 | 11.556 | 9.972 | 1.584 | −0.103 | 3.27 | 7.54% |
| P08 | 11.081 | 10.649 | 0.431 | −1.126 | 1.988 | 8.85% |
| S10 | 9.764 | 9.118 | 0.645 | −0.509 | 1.8 | 16.08% |
| U01 | 12.628 | 11.368 | 1.26 | −0.278 | 2.798 | 9.07% |
| U02 | 12.856 | 11.708 | 1.149 | −0.842 | 3.139 | 5.41% |
| U03 | 13.004 | 12.084 | 0.921 | −1.003 | 2.845 | 5.79% |
| **Meta analysis** | | | **1.031** | **0.568** | **1.494** | |

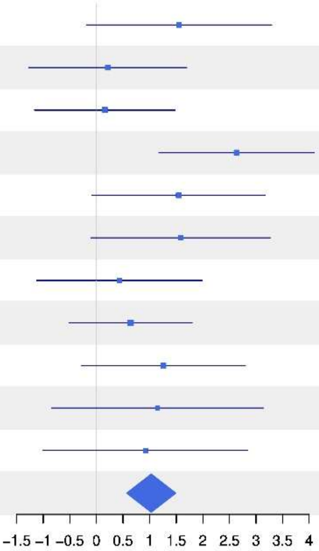


### Alpha with Coverage = 0.95

| Site | FD(q=1) Enhanced | FD(q=1) Control | Difference | LCL | UCL | W(fixed) |
|---|---|---|---|---|---|---|
| B04 | 4.501 | 4.401 | 0.1 | −0.1 | 0.301 | 8.51% |
| B05 | 5.149 | 6.203 | −1.054 | −1.305 | −0.802 | 5.42% |
| B06 | 4.05 | 3.708 | 0.343 | 0.188 | 0.497 | 14.4% |
| B07 | 3.585 | 2.768 | 0.817 | 0.696 | 0.938 | 23.47% |
| H09 | 5.408 | 4.834 | 0.573 | 0.332 | 0.815 | 5.88% |
| L11 | 6.205 | 4.098 | 2.106 | 1.803 | 2.41 | 3.72% |
| P08 | 4.062 | 4.921 | −0.859 | −1.022 | −0.696 | 12.9% |
| S10 | 4.837 | 4.729 | 0.108 | −0.1 | 0.317 | 7.88% |
| U01 | 6.787 | 4.568 | 2.219 | 2.012 | 2.426 | 8.01% |
| U02 | 7.069 | 5.2 | 1.869 | 1.607 | 2.132 | 4.97% |
| U03 | 6.707 | 6.594 | 0.112 | −0.154 | 0.379 | 4.84% |
| **Meta analysis** | | | **0.478** | **0.42** | **0.537** | |

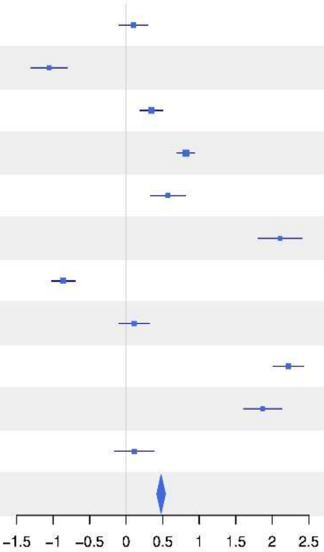

### Alpha with Coverage = 0.95

| Site | FD(q=2) Enhanced | FD(q=2) Control | Difference | LCL | UCL | W(fixed) |
|---|---|---|---|---|---|---|
| B04 | 3.193 | 3.278 | −0.085 | −0.207 | 0.037 | 9.82% |
| B05 | 3.584 | 4.589 | −1.005 | −1.184 | −0.826 | 4.57% |
| B06 | 2.986 | 2.612 | 0.374 | 0.279 | 0.468 | 16.48% |
| B07 | 2.399 | 1.723 | 0.676 | 0.606 | 0.746 | 29.7% |
| H09 | 4.193 | 3.666 | 0.527 | 0.364 | 0.69 | 5.52% |
| L11 | 4.167 | 2.849 | 1.317 | 1.078 | 1.557 | 2.56% |
| P08 | 1.625 | 3.437 | −1.812 | −1.931 | −1.694 | 10.41% |
| S10 | 3.556 | 3.472 | 0.084 | −0.072 | 0.24 | 6.06% |
| U01 | 4.761 | 2.508 | 2.253 | 2.1 | 2.407 | 6.26% |
| U02 | 4.914 | 3.015 | 1.899 | 1.72 | 2.077 | 4.61% |
| U03 | 4.566 | 4.661 | −0.095 | −0.286 | 0.096 | 4.01% |
| **Meta analysis** | | | **0.312** | **0.274** | **0.351** | |

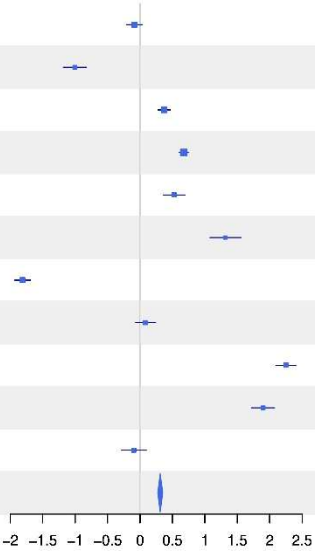


### Gamma with Coverage = 0.95

| Site | FD(q=0) Enhanced | FD(q=0) Control | Difference | LCL | UCL | W(fixed) |
|---|---|---|---|---|---|---|
| B04 | 11.544 | 9.808 | 1.736 | −5.793 | 9.266 | 6.43% |
| B05 | 12.736 | 12.345 | 0.392 | −6.729 | 7.512 | 7.19% |
| B06 | 10.323 | 9.688 | 0.635 | −4.279 | 5.549 | 15.1% |
| B07 | 11.105 | 8.412 | 2.693 | −1.197 | 6.582 | 24.1% |
| H09 | 11.048 | 9.619 | 1.429 | −5.803 | 8.662 | 6.97% |
| L11 | 13.639 | 9.874 | 3.764 | −5.014 | 12.543 | 4.73% |
| P08 | 12.116 | 10.849 | 1.267 | −5.431 | 7.966 | 8.13% |
| S10 | 10.43 | 9.831 | 0.599 | −4.197 | 5.396 | 15.85% |
| U01 | 15.827 | 13.513 | 2.313 | −5.652 | 10.278 | 5.75% |
| U02 | 16.854 | 13.346 | 3.508 | −8.158 | 15.174 | 2.68% |
| U03 | 17.03 | 13.634 | 3.396 | −7.519 | 14.31 | 3.06% |
| **Meta analysis** | | | **1.691** | **−0.218** | **3.601** | |

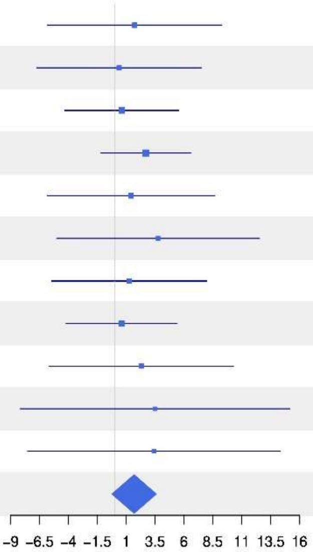

**Gamma with Coverage = 0.95**

| Site | FD(q=1) Enhanced | FD(q=1) Control | Difference | LCL | UCL | W(fixed) |
|---|---|---|---|---|---|---|
| B04 | 4.78 | 4.761 | 0.019 | −0.233 | 0.271 | 9.25% |
| B05 | 5.5 | 6.806 | −1.306 | −1.652 | −0.96 | 4.93% |
| B06 | 4.654 | 3.824 | 0.829 | 0.647 | 1.012 | 17.68% |
| B07 | 3.833 | 3.64 | 0.194 | 0.048 | 0.34 | 27.6% |
| H09 | 5.699 | 5.363 | 0.336 | 0.056 | 0.615 | 7.53% |
| L11 | 6.923 | 4.33 | 2.593 | 2.137 | 3.05 | 2.82% |
| P08 | 5.99 | 5.39 | 0.6 | 0.36 | 0.839 | 10.24% |
| S10 | 5.225 | 5.242 | −0.016 | −0.281 | 0.248 | 8.41% |
| U01 | 8.254 | 6.455 | 1.799 | 1.455 | 2.143 | 4.99% |
| U02 | 9.017 | 6.999 | 2.018 | 1.59 | 2.446 | 3.22% |
| U03 | 8.546 | 7.77 | 0.776 | 0.356 | 1.197 | 3.33% |
| **Meta analysis** | | | **0.476** | **0.4** | **0.553** | |

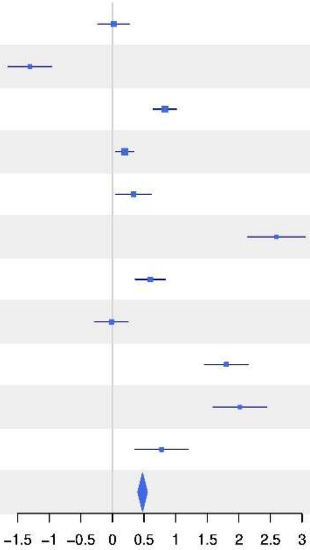


**Gamma with Coverage = 0.95**

| Site | FD(q=2) Enhanced | FD(q=2) Control | Difference | LCL | UCL | W(fixed) |
|---|---|---|---|---|---|---|
| B04 | 3.436 | 3.592 | −0.156 | −0.29 | −0.022 | 11.29% |
| B05 | 3.952 | 5.497 | −1.545 | −1.712 | −1.378 | 7.31% |
| B06 | 3.531 | 2.728 | 0.803 | 0.704 | 0.902 | 20.82% |
| B07 | 2.616 | 2.735 | −0.119 | −0.21 | −0.028 | 24.51% |
| H09 | 4.539 | 4.259 | 0.28 | 0.108 | 0.451 | 6.92% |
| L11 | 4.726 | 3.14 | 1.586 | 1.284 | 1.888 | 2.23% |
| P08 | 4.039 | 3.898 | 0.141 | −0.035 | 0.316 | 6.62% |
| S10 | 3.976 | 4.005 | −0.03 | −0.199 | 0.14 | 7.06% |
| U01 | 6.119 | 4.825 | 1.294 | 1.086 | 1.502 | 4.7% |
| U02 | 6.862 | 5.356 | 1.506 | 1.297 | 1.715 | 4.67% |
| U03 | 6.328 | 6.076 | 0.252 | 0.023 | 0.481 | 3.87% |
| **Meta analysis** | | | **0.21** | **0.165** | **0.256** | |

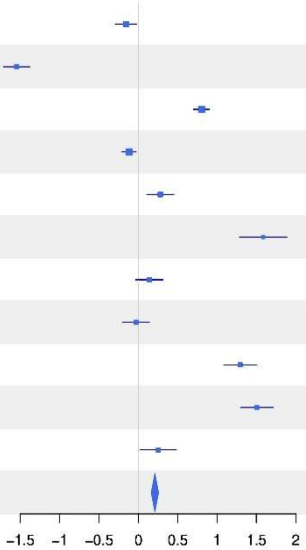

#### 1−S with Coverage = 0.95

| Site | FD(q=0) Enhanced | FD(q=0) Control | Difference | LCL | UCL | W(fixed) |
|---|---|---|---|---|---|---|
| B04 | −0.005 | 0.003 | −0.008 | −0.092 | 0.075 | 5.59% |
| B05 | −0.002 | 0.008 | −0.01 | −0.093 | 0.072 | 5.69% |
| B06 | 0.004 | 0.004 | 0 | −0.083 | 0.084 | 5.58% |
| B07 | 0.003 | 0.016 | −0.013 | −0.103 | 0.076 | 4.83% |
| H09 | 0.005 | 0.002 | 0.003 | −0.092 | 0.098 | 4.33% |
| L11 | 0.011 | −0.011 | 0.022 | −0.089 | 0.133 | 3.17% |
| P08 | 0.008 | −0.014 | 0.023 | −0.053 | 0.099 | 6.7% |
| S10 | 0.007 | 0.013 | −0.006 | −0.092 | 0.08 | 5.25% |
| U01 | 0.024 | 0.025 | −0.001 | −0.049 | 0.047 | 16.95% |
| U02 | 0.028 | 0.013 | 0.015 | −0.025 | 0.056 | 23.62% |
| U03 | 0.023 | 0.007 | 0.016 | −0.03 | 0.062 | 18.29% |
| **Meta analysis** | | | **0.007** | **−0.013** | **0.026** | |

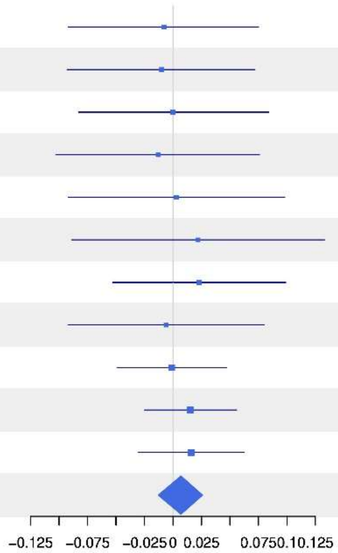


#### 1−S with Coverage = 0.95

| Site | FD(q=1) Enhanced | FD(q=1) Control | Difference | LCL | UCL | W(fixed) |
|---|---|---|---|---|---|---|
| B04 | 0.046 | 0.055 | −0.009 | −0.026 | 0.009 | 7.95% |
| B05 | 0.047 | 0.057 | −0.01 | −0.029 | 0.008 | 7.02% |
| B06 | 0.098 | 0.026 | 0.072 | 0.059 | 0.085 | 14.67% |
| B07 | 0.05 | 0.226 | −0.176 | −0.189 | −0.163 | 15.39% |
| H09 | 0.027 | 0.069 | −0.042 | −0.062 | −0.022 | 6% |
| L11 | 0.052 | 0.043 | 0.008 | −0.016 | 0.032 | 4.19% |
| P08 | 0.289 | 0.056 | 0.234 | 0.22 | 0.247 | 13.79% |
| S10 | 0.051 | 0.068 | −0.017 | −0.038 | 0.004 | 5.45% |
| U01 | 0.079 | 0.222 | −0.142 | −0.159 | −0.126 | 9% |
| U02 | 0.109 | 0.191 | −0.082 | −0.099 | −0.065 | 8.33% |
| U03 | 0.101 | 0.085 | 0.017 | 0 | 0.034 | 8.22% |
| **Meta analysis** | | | **−0.007** | **−0.012** | **−0.002** | |

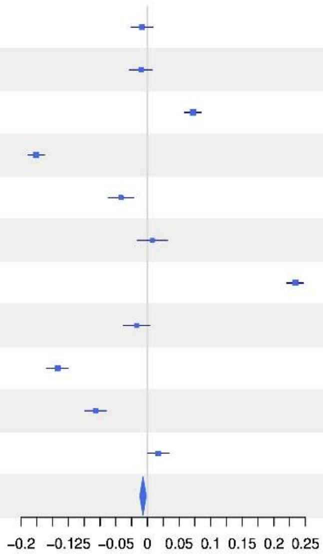

### 1−S with Coverage = 0.95

| Site | FD(q=2) Enhanced | FD(q=2) Control | Difference | LCL | UCL | W(fixed) |
|---|---|---|---|---|---|---|
| B04 | 0.073 | 0.077 | −0.004 | −0.02 | 0.012 | 14.02% |
| B05 | 0.085 | 0.122 | −0.037 | −0.063 | −0.011 | 5.44% |
| B06 | 0.14 | 0.044 | 0.096 | 0.078 | 0.114 | 11.28% |
| B07 | 0.081 | 0.393 | −0.312 | −0.329 | −0.295 | 12.29% |
| H09 | 0.043 | 0.11 | −0.067 | −0.091 | −0.043 | 6.24% |
| L11 | 0.079 | 0.084 | −0.004 | −0.032 | 0.024 | 4.65% |
| P08 | 0.621 | 0.097 | 0.524 | 0.507 | 0.54 | 12.96% |
| S10 | 0.088 | 0.107 | −0.019 | −0.041 | 0.002 | 7.82% |
| U01 | 0.136 | 0.441 | −0.305 | −0.324 | −0.286 | 10.14% |
| U02 | 0.189 | 0.372 | −0.184 | −0.204 | −0.163 | 8.57% |
| U03 | 0.179 | 0.153 | 0.026 | 0.003 | 0.05 | 6.61% |
| **Meta analysis** | | | **−0.013** | **−0.019** | **−0.007** | |

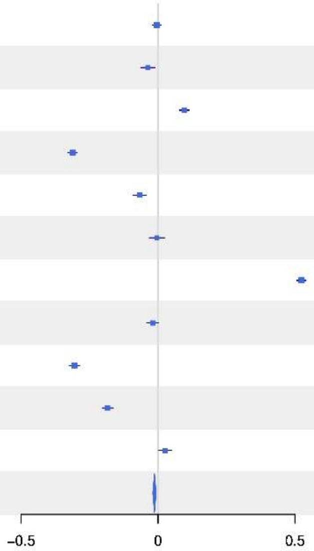

Table S1: Results of linear mixed interaction models of the beta-deviation, i.e. the difference between beta diversity of observed vs. null-model data (with randomized distribution of individuals across patches in each site) for taxonomic (TD), functional (FD), and phylogenetic (PD) beta diversity in combination with Hill numbers q=0, 1, 2 (rare, common, and dominant species). As predictors we used the structural difference between patches (DistTreat), the difference in space (DistXY), and the difference in abiotic conditions (DistPlant) extracted from vegetation indicator values in interaction with manipulated deadwood and canopy structure (TreatC vs. TreatE, i.e. control vs. enhanced), as well as Site as random factor.

| Diversity facet | TD | TD | TD | FD | FD | FD | PD | PD | PD |
|---|---|---|---|---|---|---|---|---|---|
| Hill number q | 0 | 1 | 2 | 0 | 1 | 2 | 0 | 1 | 2 |
| TreatE | 0.975 | -1.237 | **-4.808** | -0.168 | **-4.900** | **-5.947** | 1.830 | **-4.262** | **-5.362** |
| DistTreat | **3.753** | **2.731** | 0.204 | -0.111 | -1.010 | -1.308 | **2.430** | -0.410 | -0.853 |
| DistXY | 1.169 | **5.816** | **4.578** | 0.725 | **3.268** | **3.257** | 1.582 | **4.501** | **4.133** |
| DistPlant | 0.085 | -1.501 | **-2.039** | 1.127 | **-3.717** | **-3.571** | 0.164 | **-3.665** | **-4.020** |
| TreatE:DistTreat | -0.727 | **4.298** | **5.276** | **2.147** | **2.496** | **3.032** | -1.245 | **2.467** | **2.102** |
| TreatE:DistXY | -0.187 | **-2.218** | -1.199 | -0.502 | -0.473 | -0.121 | -0.217 | -1.535 | -1.023 |
| TreatE:DistPlant | 0.468 | **2.449** | **4.745** | **-2.294** | **4.861** | **5.388** | 1.093 | **5.353** | **5.666** |

Figure S28: Standardized effect sizes of mean pairwise distances in functional β-diversity of control and enhanced patches in response to treatment categories for rare (q=0), common (q=1), and dominant (q=2) species. P-values for intercept and ESBC treatment indicate if SES values in general significantly differ from 0 and if ESBC differs from Control in a generalized mixed effect model of SES in response to treatment. Percentage values represent the fraction of replicates above (red) or below (below) the vertical 1.96 lines.

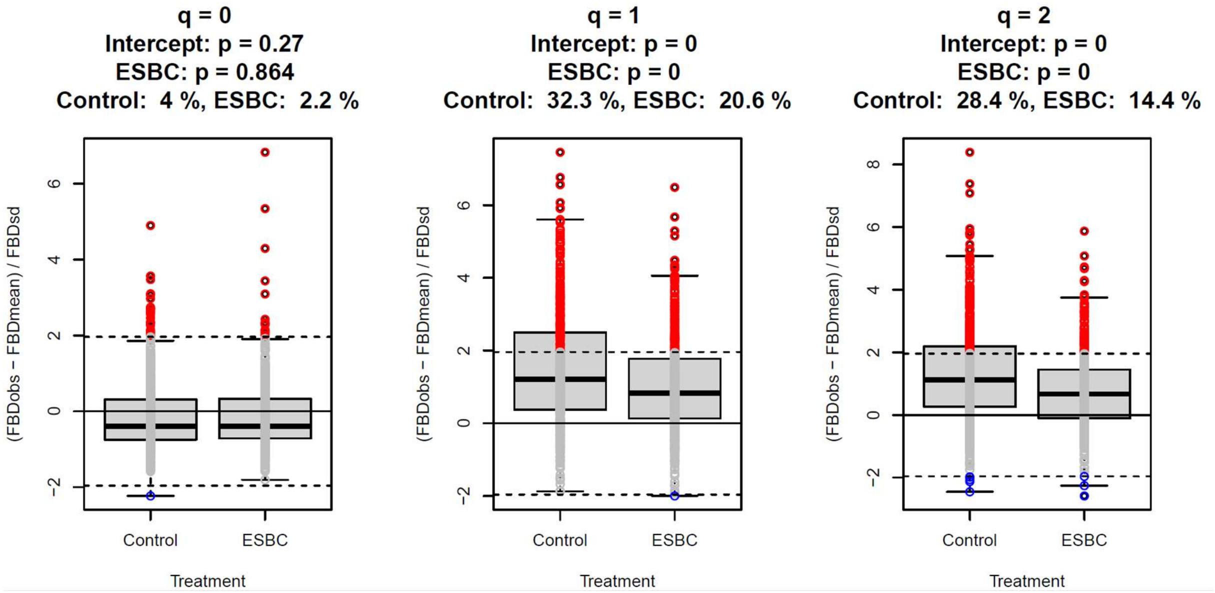
